## Supplemental Figures for "A secreted protease-like protein in *Zymoseptoria tritici* is responsible for avirulence on *Stb9* resistance gene in wheat"

### Supporting information

**Table S1.** Mean disease phenotypes measured among wild-types, mutant and chimeric strains of AvrStb9 alleles.

**Table S2.** Summary statistics of GWA analysis on Stb9 resistance in the BW panel. Only SNPs above the Bonferroni-corrected threshold are presented.

**Table S3.** List of the top 10 protein tertiary structures with the highest similarity to AvrStb9 protein structure (from DALI webserver; accessed in january 2023).

**Table S4.** List of the pairwise structure alignment results.

**Table S5.** List of AvrStb9 selected orthologs/paralogs identified by blastp search using OrthoMCL database.

**Table S6.** Model test and parameter estimates of diversifying selection with codeml based on the total *AvrStb9* data set.

**Table S7.** List of primers used in this study.

**Figure S1.** Density distribution of percentage of leaf area covered by pycnidia (PLACP) in the fungal population.

**Figure S2.** Population structure and relatedness in the fungal population used for GWAS analysis.

**Figure S3.** Manhattan plots of PLACN and PLACP on the wheat cultivar 'Soissons'.

**Figure S4.** Improved *AvrStb9* gene model using RNA-seq reads mapped to the reference genome IPO323.

**Figure S5.** Wild-type, mutant and chimeric strains phenotype (PLACP) on cultivars carrying or not the resistance gene *Stb9* and the susceptible cultivar Taichung-29.

**Figure S6.** Distribution of the relative AUDPC sporulating leaf area in the bread wheat population and lead GWAS SNP phenotypes.

**Figure S7.** *AvrStb9* paralogs and their expression profile.

**Figure S8.** Disease scale used for the visual assessment of percent of leaf area covered by necrosis (PLACN) and by pycnidia (PLACP) after inoculation of wheat leaves with the phytopathogenic fungus *Zymoseptoria tritici*.

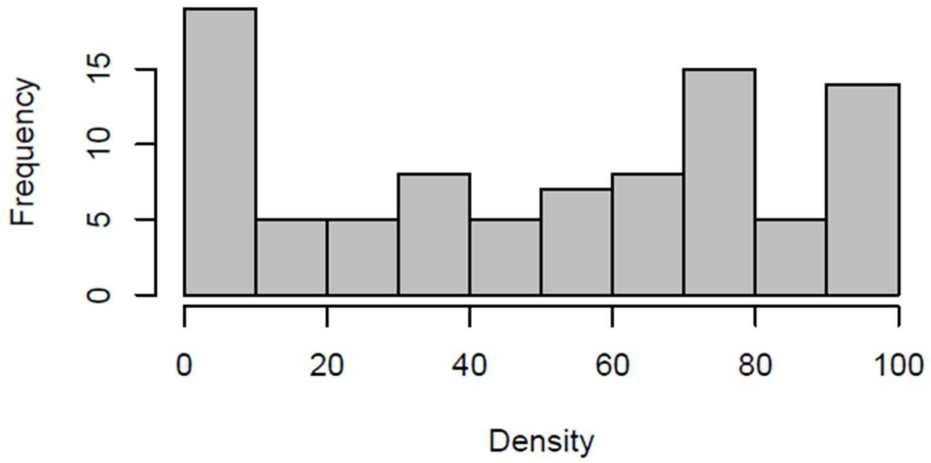


**Figure S1.** Density distribution of percentage of leaf area covered by pycnidia (PLACP) measured at 21 days post inoculation (dpi) on the wheat cultivar 'Soissons' in the *Z. tritici* fungal population.

**(a)** **(b)**

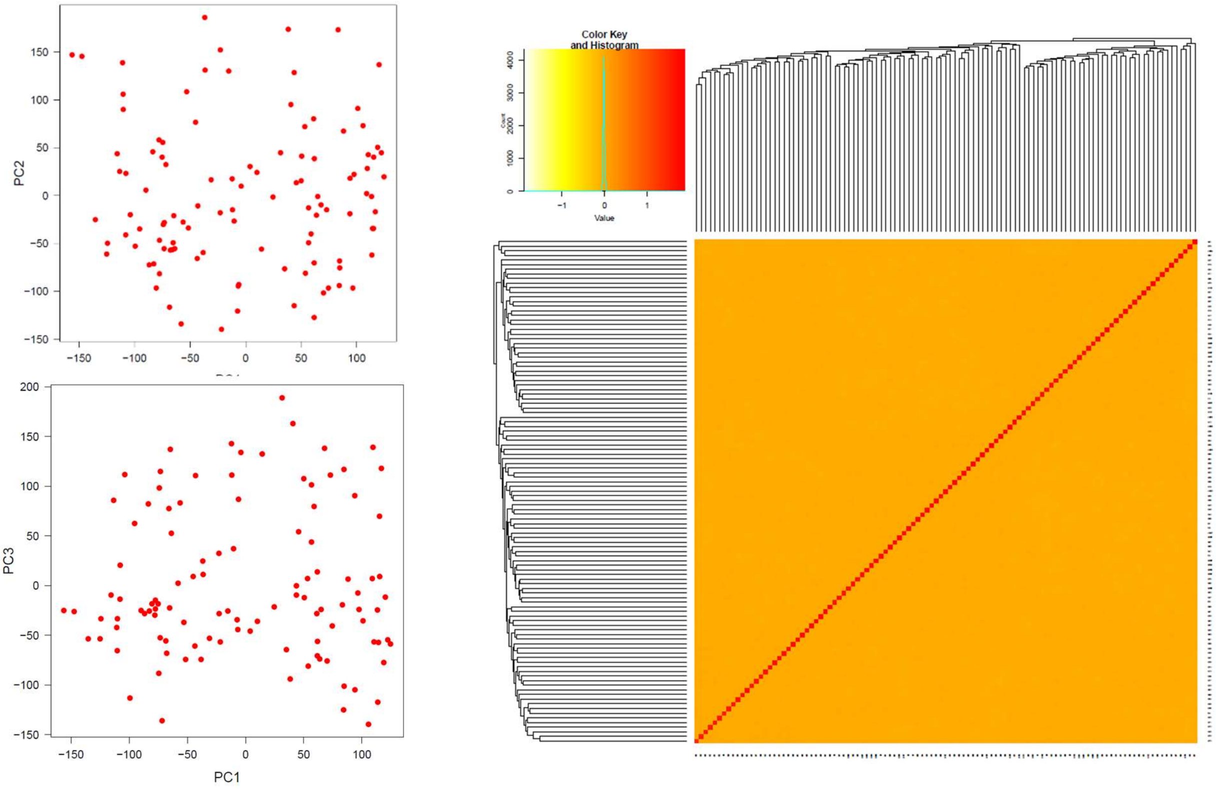


**Figure S2.** Population structure and relatedness in the fungal population used for GWAS analysis. (a) Principal component analysis using full SNP data. PC1, PC2 and PC3 explained 1.30%, 1.12% an 0.69% of the phenotypic variation, respectively. (b) Heatmap of the kinship matrix using VanRaden method (VanRaden, 2008).

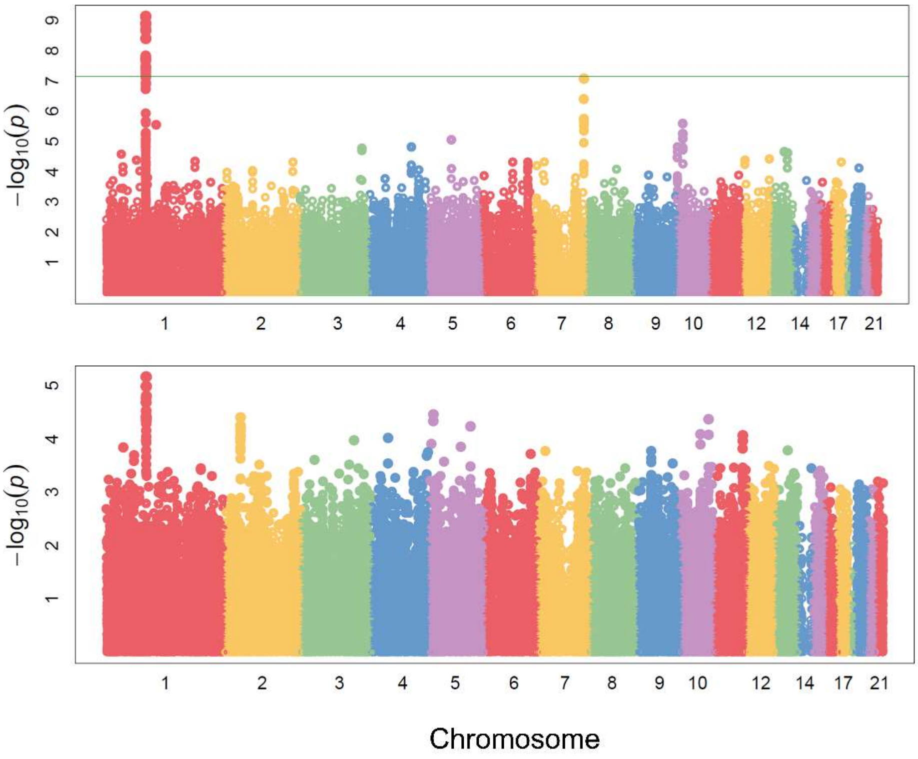


**Figure S3.** Manhattan plots of PLACN (top) and PLACP (bottom) on the wheat cultivar 'Soissons'. The horizontal line indicates the genome-wide significance threshold (Bonferroni correction at $\alpha<0.05$).

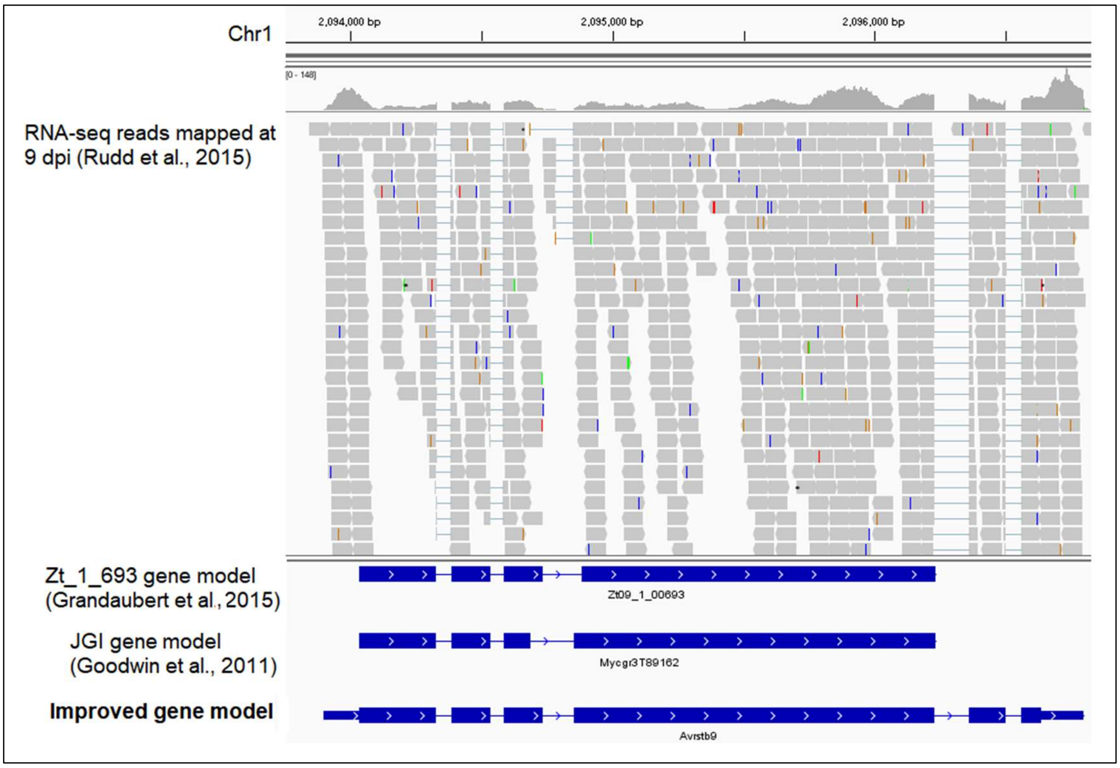


**Figure S4.** Improved *AvrStb9* gene model using RNA-seq reads at 9 dpi mapped to the reference genome IPO323.

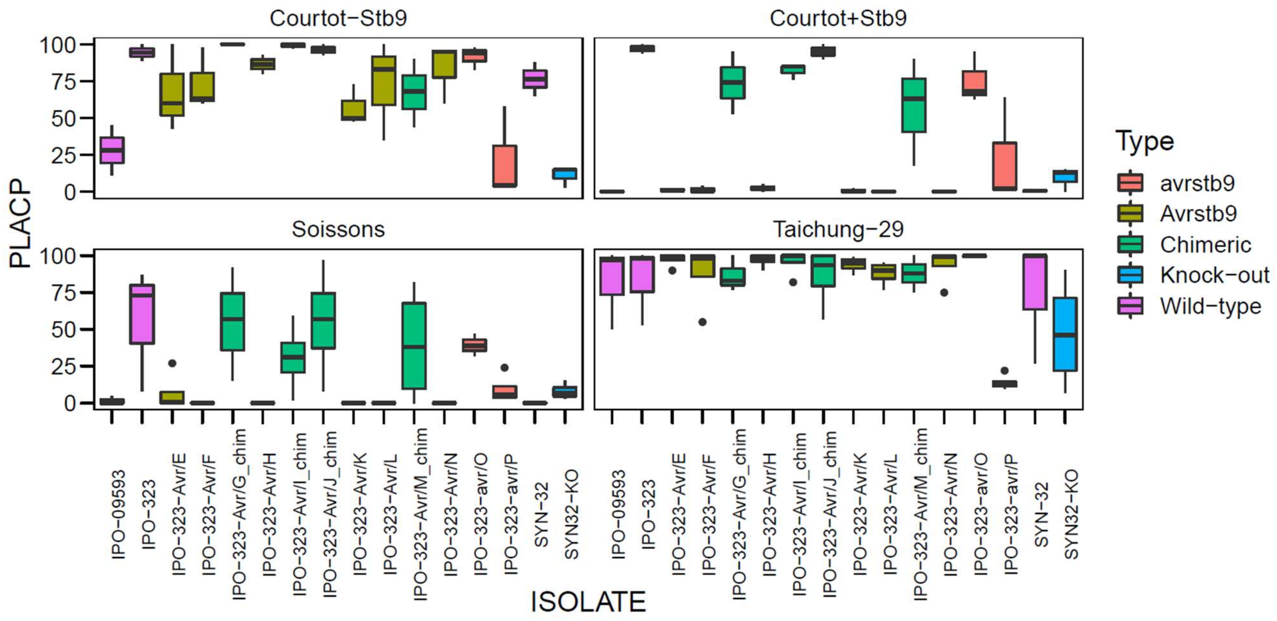


**Figure S5.** PLACP of wild-type, mutant, chimeric strains and the single knock-out mutant phenotyped on cultivars carrying or not the resistance gene *Stb9* and the susceptible cultivar 'Taichung-29'.

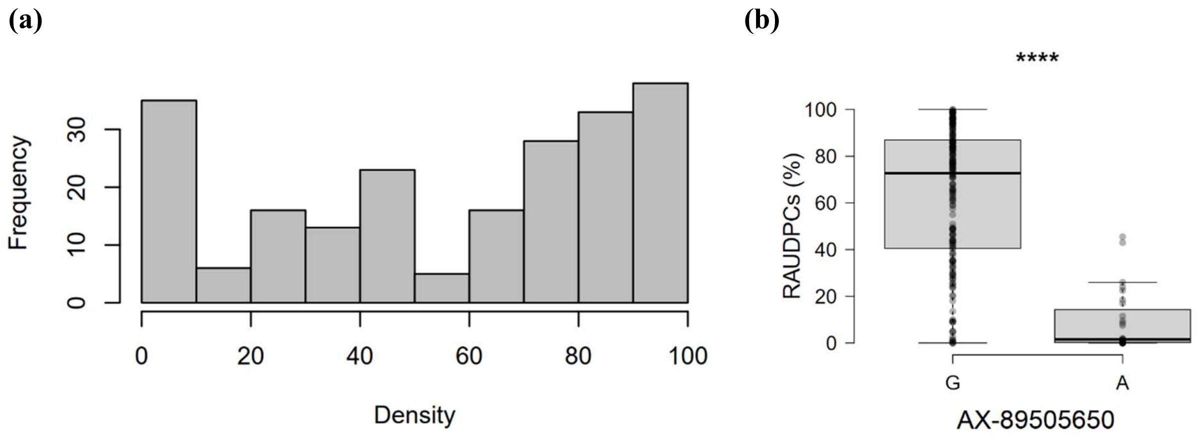


**Figure S6.** Distribution of the relative AUDPC sporulating leaf (RAUDPCs) area in the panel of bread wheat cultivars (a) and the lead SNP of *Stb9* locus mapped using GWAS in the same panel (b).

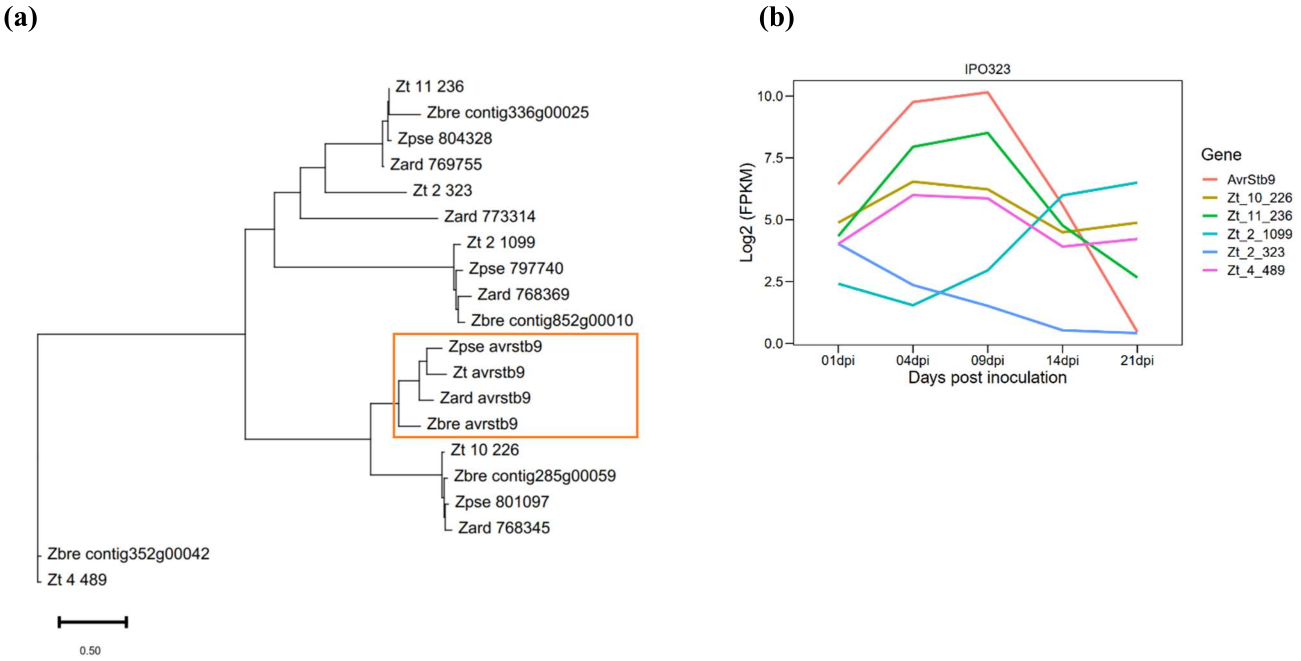


**Figure S7.** *AvrStb9* homologs and their expression profile. (a) A phylogenetic tree of *AvrStb9* paralogs and orthologs from *Z. tritici* and its relative species (*Zt=Z.tritici; Zpse=Z. pseudotritici; Zbre=Z.brevis; Zard= Z. ardabilae*). (b) Gene expression profile of *AvrStb9* paralogs using IPO323 RNAseq data (Rudd *et al.*, 2015).

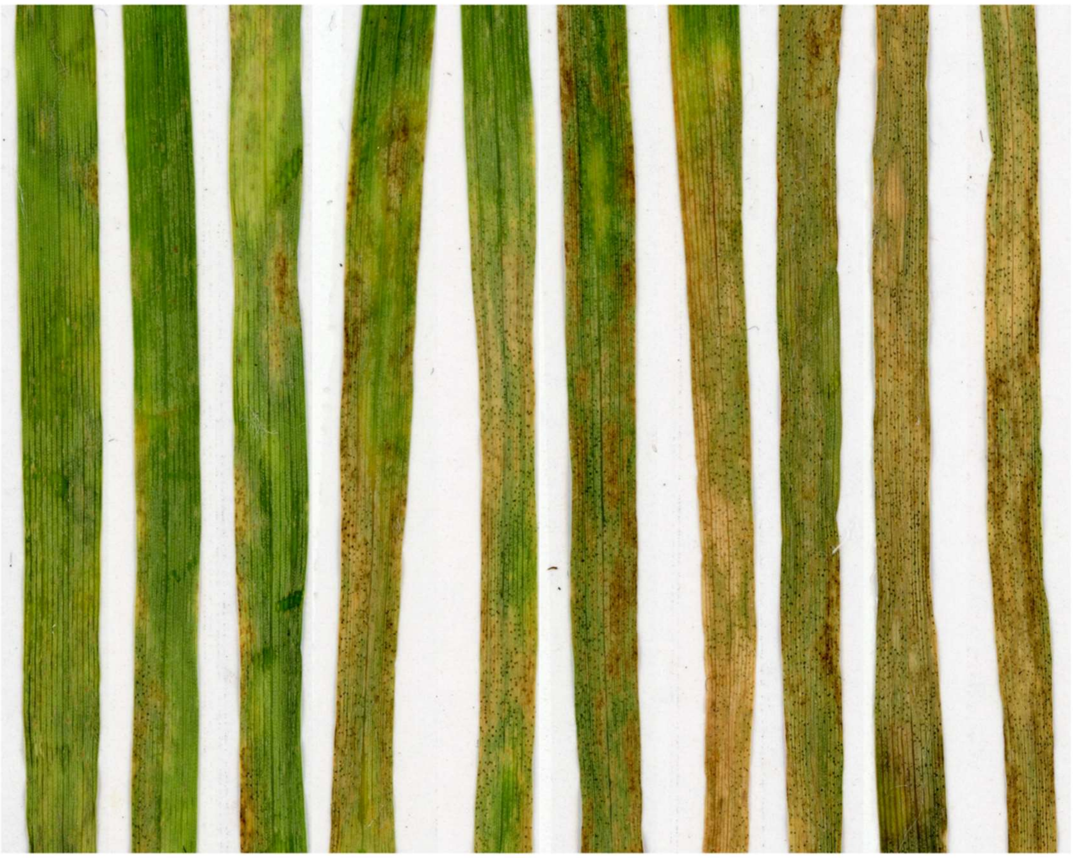

| | | | | | | | | | | |
|---|---|---|---|---|---|---|---|---|---|---|
| PLACN | 0% | 10% | 20% | 40% | 50% | 80% | 90% | 100% | 100% | 100% |
| PLACP | 0% | 5% | 5% | 35% | 50% | 65% | 70% | 80% | 85% | 100% |

**Figure S8.** Disease scale used for the visual assessment of percent of leaf area covered by necrosis (PLACN) and by pycnidia (PLACP) after inoculation of wheat leaves with the phytopathogenic fungus *Zymoseptoria tritici*.
